# Molecular Dosimetry of DNA Adducts in Mice Exposed to Ethylene Oxide

**DOI:** 10.64898/2026.03.25.714191

**Authors:** Chih-Wei Liu, Jingya Peng, Jiahao Feng, Haoduo Zhao, Xueying Wang, B. Bhaskar Gollapudi, Abby A. Li, James S. Bus, Kun Lu

## Abstract

Ethylene oxide (EtO) is a highly reactive industrial chemical and classified as a known human carcinogen with a putative mutagenic mode of action (MOA). Its genotoxic potential is primarily mediated through alkylation of DNA, resulting in the formation of the mutagenic adduct O^6^-(2-hydroxyethyl)-2’-deoxyguanosine (O^6^-HE-dG). The N7-(2-hydroxyethyl)guanine (N7-HE-G) adduct is formed in greater abundance and is generally considered to be non-mutagenic. However, dose-response relationships of these DNA adducts, particularly at low inhalation exposure levels (i. e., below 3 ppm), remain unknown. These data are necessary to inform the biological plausibility of different statistical dose-response models that have been applied to human or animal data used for cancer risk assessment. In the present study, male and female B6C3F1 mice were exposed to EtO (0, 0.05, 0.1, 0.5, 1, 50, 100, and 200 ppm) 6 hours/day for 28 consecutive days. Immediately following the last exposure, DNA was extracted from lung, liver, bone marrow, and mammary gland, and further utilized to measure DNA adduct levels using highly sensitive mass spectrometry platforms. N7-HE-G was detected in all tissues and exposure groups, showing linear dose-response relationships in the low-dose range (≤1 ppm) and increased sharply and exposure-disproportionately in the high-dose range (≥50 ppm). Despite a very low limit of detection, O^6^-HE-dG, in contrast, was not detected at exposures <50 ppm in any tissue consistent with at most a shallow linear exposure response. At higher exposures (≥50 ppm), O^6^-HE-dG exhibited a dose-response pattern of N7-HE-G. Notably the mammary gland, despite being anatomically distant from the site of inhalation, exhibited the second-highest levels of both adducts at higher doses. This study provides the first reliable quantitative dose-response evidence of DNA adducts in tumor target and non-target (liver) tissues across a wide range of EtO exposures. The two DNA adducts differ markedly in their abundance, repairability and mutagenic potential and together provide a molecular MOA dose-response framework to inform both quantitative cancer risk assessment and genotoxic hazard characterization.

## INTRODUCTION

Ethylene oxide (EtO) is a colorless, highly reactive gas with an estimated global production of around 20 million tons, ranking it among the most extensively produced organic chemicals worldwide (Ghosh and Godderis 2016). Its primary industrial application is as a chemical intermediate in the synthesis of products such as emulsifiers, detergents, solvents, plastics, textiles, and antifreeze (Eastmond et al. 2014). In addition to its industrial uses, EtO serves a critical function as a sterilizing agent for heat-sensitive medical and dental equipment and is employed as a fumigant for the decontamination of various food products, including spices (Lynch et al. 2022). Exposures can occur to workers that make or use EtO, and to the general population through its release to the environment (primarily air), and to all populations via its endogenous production from ethylene (Kirman et al. 2021; Kirman et al. 2025). A mean US total daily general population exposure has been reported as 3.5 ppb resulting from a mean of 0.2 ppb in ambient air and 3.3 ppb attributed to endogenously generated EtO. Cigarette smoke contains ethylene and EtO, and total daily exposure for approximately 1 pack/day smokers is equivalent to 16.6 ppb/day EtO (Kirman et al. 2025). Recommended occupational exposure limits have been set at 1 ppm since 1984 (ACGIH TLV 1984; OSHA PEL 1984).

The International Agency for Research on Cancer (IARC 2008), United States Environmental Protection Agency (USEPA 2016) and Texas Commission on Environmental Quality (TCEQ 2020) have classified EtO as a known or likely human carcinogen. These classifications are based on limited epidemiological evidence primarily for lymphoid and/or breast cancers. The cancer classifications were supported by evidence of rat and mouse carcinogenicity induced by chronic inhalation exposures from 10 to 100 ppm, as well as associated mode of action evidence (IARC 2008; TCEQ 2020; USEPA 2016).

As a direct-acting alkylating agent, the EtO MoA is generally attributed to reactions with nucleophilic sites on DNA, RNA, and proteins, leading to genotoxic and cancer effects at higher exposures (Eastmond et al. 2014; Gollapudi et al. 2025; Gollapudi et al. 2020; Vincent et al. 2019). Among the EtO DNA adducts, the N7-(2′-hydroxyethyl)-guanine (N7-HE-G) adduct is the most abundant, accounting for approximately 95% of all EtO-induced DNA adducts (Segerback 1990; Swenberg et al. 2011). This adduct forms through alkylation at the N7 position of guanine, which is the most nucleophilic site among DNA bases and commonly reacts with alkylating agents (Gates et al. 2004). Although N7-HE-G itself is not considered promutagenic, it is chemically unstable and prone to spontaneous depurination with formation of apurinic/apyrimidinic (AP) sites. Despite this theoretical mutagenic potential, repeated exposure to a EtO 100 ppm tumorigenic exposure did not increase AP site DNA damage (Rusyn et al. 2005; Swenberg et al. 2011). The half-life of N7-HE-G varies depending on the tissue and conditions, but it is generally short, ranging from a few hours to several days in double-stranded DNA (Margison et al. 1976). Given its short half-life, N7-HE-G accumulates to steady-state levels typically reached after 7–10 days of repeated exposure to EtO in rodents and as projected by PBPK modeling in humans (Filser and Klein 2018; Walker et al. 2000). At steady-state levels, the number of N7-HE-G adducts formed is equal to the number of adducts lost due to depurination, repair, or cell death (Pottenger et al. 2019). This predictable accumulation pattern makes the N7-HE-G adduct a useful biomarker for internal EtO dose and exposure duration, even though it is not directly mutagenic.

Although O^6^-(2’-hydroxyethyl)-2′-deoxyguanosine (O^6^-HE-dG) is a minor DNA adduct formed from EtO exposure, it is considered as a critical DNA lesion due to its strong mutagenic potential. O^6^-HE-dG adduct is potentially mispaired with thymine during DNA replication, leading to G:C→A:T transition mutations (Delaney and Essigmann 2001; Mazon et al. 2010). Unlike N7-HE-G adducts, O^6^-HE-dG adducts exhibit limited spontaneous depurination and are inefficiently repaired in the absence of dedicated repair pathways. Persistence of O^6^-HE-dG induces activation of the mismatch repair (MMR) system via recognition of O^6^-alkylguanine: thymine mispairs, leading to cycles of futile repair and subsequent induction of apoptosis through DNA damage response signaling cascades (Mazon et al. 2010). *In vivo* investigations have demonstrated that O^6^-HE-dG adducts in rat tissues attain steady-state concentrations following two weeks of EtO inhalation exposure at 300 ppm (Walker et al. 1992). The relative stability and biological impact of O^6^-HE-dG adducts implicate them as principal contributors to EtO-mediated mutagenesis and carcinogenesis.

While dose–response relationships between EtO and DNA adduct formation have been demonstrated in various *in vitro* and *in vivo* models, the precise characterization of these relationships at low inhalation exposures that are relevant to the general population remains limited. Several studies have shown a clear dose-dependent increase in N7-HE-G adduct formation at exposures above 1 ppm. Walker et al. reported significant increases in N7-HE-G in the brain, spleen, and lung of F344 rats beginning at 3 ppm of EtO exposures (Walker et al. 1990; Walker et al. 1992). They also observed that N7-HE-G adduct levels in B6C3F1 mice were approximately 2- to 3-fold lower than in the same tissues of concurrently exposed rats. In contrast, O^6^-HE-dG adducts occur at nearly 300 times lower than N7-HE-G adducts *in vivo*, limiting the ability of most studies to establish comprehensive dose–response curves for this adduct across a broad range of exposures. Notably, Walker et al. only observed a significant increase in O^6^-HE-dG adducts at EtO concentrations of 300 ppm (Walker et al. 1992). Although elevated mutation frequencies in bone marrow and increased breast cancer risk have been linked to EtO exposure (USEPA 2016), specific dose–response data for N7-HE-G and O^6^-HE-dG adducts in these organs remain lacking.

To address these gaps, our study aims to evaluate the dose–response relationships of both N7-HE-G and O^6^-HE-dG adducts in male and female B6C3F1 mice, the strain used in the EtO cancer bioassay (National Toxicology Program 1987). Mice aged 9–12 weeks were exposed to EtO via whole-body inhalation for 6 hours each day over a 28-day period. This exposure period was selected because both adducts reach steady state by 4 weeks and therefore provide adduct dose-response data needed to evaluate the biological plausibility of differing statistical models of cancer risk applied to human studies. Eight exposure groups were tested: 0 ppm (air control), 0.05 ppm, 0.1 ppm, 0.5 ppm, 1 ppm, 50 ppm, 100 ppm, and 200 ppm.

## MATERIALS AND METHODS

### Chemicals and Materials

Unless otherwise specified, all reagents and chemicals used in this study were purchased from Sigma Aldrich (St. Louis, MO). EtO (CAS Number 75-21-9; purity 99.99%) was provided by Balchem Corporation, (Green Pond, SC). The Optima LC-MS grade methanol (MeOH), acetonitrile (ACN), water, isopropyl alcohol (IPA) and formic acid (FA) were all purchased from Thermo Fisher Scientific (Rockford, IL). NanoSep Centrifugal Devices (MWCO 3K) and stainless steel beads (5 mm) were obtained from Pall Life Sciences (Port Washington, NY) and QIAGEN (Germantown, MD), respectively. NucleoBond AXG 20 columns and NucleoBond buffer kits were purchased from Macherey-Nagel (Bethlehem, PA). Proteinase K was obtained from VWR International, LLC (Atlanta, GA). Breathe-easier breathable tube membranes for microtubes were purchased from Sigma Aldrich (product no. Z743501). Synthetic standards, N7-(2-hydroxyethyl)guanine (N7-HE-G, TRC-H942200) and O^6^-(2-hydroxyethyl)-2′-deoxyguanosine (O^6^-HE-dG, TRC-H942020), and stable isotope labeled internal standard (IST), N7-(2-hydroxyethyl)guanine-d4 (N7-HE-G-d4, TRC-H942202) and O^6^-(2-hydroxyethyl)-2′-deoxyguanosine-d4 (O^6^-HE-dG-d4, TRC-H942022) were purchased from LGC Standards (Manchester, NH).

### Mouse Exposure Experiment with EtO

Mice were exposed via whole-body inhalation to filtered air (Group 1, G1), or EtO at concentrations of 0.05 (G2), 0.1 (G3), 0.5 (G4), 1 (G5), 50 (G6), 100 (G7), and 200 (G8) ppm for 6 h per day over 28 consecutive days. Exposures were conducted in 1000-L stainless-steel and glass whole-body exposure chambers. Due to technical issues in generating steady-state concentrations of G2 and G3 during the initial phase (Phase 1), a second phase (Phase 2) was performed, including only G1, G2, G3, and G6 to ensure exposure accuracy, as described in detail previously (Liu et al. submitted). While DNA adducts were quantified for all animals, G2 and G3 data from Phase 1 were excluded from the final dose-response analysis to maintain data integrity. For details regarding the inhalation exposure system and environmental monitoring, refer to the Supporting Information.

Immediately (within 2 hours) after the final exposure, the mice were sacrificed and tissue samples including lung, liver, bone marrow and mammary gland were collected and properly stored at –80 °C before further adduct analysis. Blood samples were also collected for dose-response analyses of EtO systemic exposure, measured as N-(2-hydroxyethyl)-L-valine (HE-V) accumulation (Liu et al. submitted), and for genotoxicity endpoints, including micronucleus and *Pig-a* assays (Gollapudi 2023). Comparative analysis of the common exposure groups (G1 and G6) across Phase I and II revealed consistent dose-dependent response trends with no significant interaction between experimental phases, justifying the integration of data from both phases for comprehensive dose-response modeling (see Supporting Information for detailed statistical validation).

### DNA Extraction

The experimental procedures for genomic DNA extraction were previously described (Hsiao et al. 2022; Liu et al. 2021). In brief, tissue samples (∼40 mg for lung and liver samples; total collected samples for bone marrow and mammary gland tissues) were homogenized in G2 solution from a NucleoBond buffer kit by a stainless steel bead via the mechanical disruption of TissueLyzer (QIAGEN) for 10 min (50 Hz). Those homogenized samples in G2 solution were further used for DNA purification according to the manufacturer’s instruction for NucleoBond AXG 20 column sample preparation. Purified DNA was reconstituted in 100 μL of water and further quantified by a Nanodrop One spectrophotometer (dsDNA/slope mode, Thermo Fisher Scientific). Extracted tissue DNA samples were further aliquoted for N7-HE-G and O^6^-HE-dG adduct analysis. For N7-HE-G, up to 20 μg of DNA was used for adduct release. However, due to limited DNA yield in bone marrow and mammary gland samples, only 5, 10, or 15 μg of DNA was used for some specimens from groups 6–8. Similarly, for O^6^-HE-dG, up to 20 μg of DNA was utilized depending on the amount of DNA remaining.

### N7-HE-G Adduct Release and Purification

DNA solution prepared in 90 μL of water was spiked with 10 μL of N7-HE-G-d4 IST (2 nM in water). N7-HE-G was released from DNA by neutral thermal hydrolysis. DNA solution was incubated in hot water bath (95 °C) for 45 min. The hydrolyzed DNA was filtered with a NanoSep 3 kDa filter (prewashed with water four times) at 8000 g for 20 min and the filtrate was further used for HPLC purification of N7-HE-G adduct.

The filtrate (80 μL) was injected into an Agilent 1200 Series UV HPLC fraction collection system for purification of target DNA adducts. Analytes were separated by reversed-phase liquid chromatography with an Atlantis C18 T3 column (150 × 4.6 mm, 3 μm, Waters). Detection wavelength and column temperature were set at 254 nm and 30 °C, respectively. Mobile phases were water with 10 mM ammonium acetate (A) and methanol (B). HPLC gradient conditions were shown in **Supplementary Table 1**. The target fraction was collected, completely dried under the breathable tube membranes in the SpeedVac Vacuum concentrator before further reconstitution in 20 μL (for G1–G5 samples) or 200 μL (for G6–G8 samples) of water for Q Exactive HF MS and TSQ Quantis QqQ MS analysis, respectively.

### LC-ESI-MS/MS Analysis of N7-HE-G Adduct

ForG1–G5 samples, nanoLC-ESI-MS/MS analysis was conducted with an UltiMate 3000 RSLCnano system coupled to a Q Exactive HF Hybrid Quadrupole-Orbitrap mass spectrometer through an EASY-Spray ion source for nanoelectrospray ionization (Thermo Fisher Scientific). The N7-HE-G fraction was separated on a PepMap C18 analytical column (2 μm particle size, 25 cm x 75 μm i.d., catalog no. ES902). Samples (6 μL) were loaded into an Acclaim PepMap C18 trapping column (3 μm particle size, 15 cm x 75 μm i.d., catalog no. 164535) at a flow rate of 5 μL/min for 2.75 min using 0.1% FA in water as a loading solvent. After 2.75 min trapping time, trapped analytes were eluted to an analytical column for separation. A binary solvent system consisting of 0.1% FA in water (solvent A) and 0.1% FA in ACN (solvent B) was used for LC separation at a flow rate of 300 nL/min. Targeted parallel reaction monitoring (PRM) MS/MS data were acquired in the positive profile mode. For the detection of N7-HE-G, an inclusion list was comprised of *m*/*z* 196.0829 (N7-HE-G) and *m*/*z* 200.1080 (N7-HE-G-d4). LC gradient conditions and MS-PRM parameters are shown in **Supplementary Table 2**.

For G6–G8 samples, on the other hand, LC-MS/MS analysis was performed using a Vanquish UHPLC system coupled to a TSQ Quantis triple quadrupole (QqQ) mass spectrometer through a Heated Electrospray Ionization probe (HESI, Thermo Fisher Scientific). The HESI ion source condition was used with default settings for a flow rate at 200 μL/min. LC separation was achieved by using an ACQUITY UPLC HSS T3 column (1.8 μm particle, 150 x 2.1 mm i.d.). A binary solvent system consisting of 0.1% FA in water (solvent A) and 0.1% FA in ACN (solvent B) was used for LC separation. Targeted selected reaction monitoring (SRM) MS/MS data were acquired in the positive centroid mode, transitions *m*/*z* 196.1 > *m*/*z* 152 and *m*/*z* 200.1 > *m*/*z* 152 for N7-HE-G and N7-HE-G-d4, respectively. LC gradient conditions and MS-SRM parameters are shown in **Supplementary Table 2**.

### O^6^-HE-dG Adduct Release, Purification, and Analysis

The O^6^-HE-dG adduct was released from DNA by enzymatic digestion. DNA solution (100 μL) was added with 200 μL of 50 mM sodium phosphate/20 mM MgCl_2_ buffer (pH 7.2) along with 10 μL of the O^6^-HE-dG-d4 IST (1 nM in water) before digestion by DNase I, alkaline phosphatase, and phosphodiesterase for 1 h at 37 °C with gentle shaking (Hsiao et al. 2022; Liu et al. 2021). The final enzymatic reaction volume was 410 μL. Following digestion, hydrolyzed DNA was filtered with a NanoSep 3 kDa filter (prewashed with water four times) at 8000 g for 50 min to remove enzymes prior to HPLC purification of O^6^-HE-dG adduct. The filtrate (390 μL) was injected into the aforementioned HPLC fraction collection system for adduct purification using the same C18 column and solvent systems. HPLC gradient conditions are shown in **Supplementary Table 1**. The target fraction was collected and completely dried before further reconstitution in 15 μL of 0.1% FA for Q Exactive HF MS analysis. Additionally, the amount of digested dG in each sample was quantitated by UV peak area based on each freshly prepared dG calibration curve to estimate the dG amount in each sample loaded into the HPLC column for adduct purification. The measured dG amount was further utilized to normalize the O^6^-HE-dG adduct numbers.

The nanoLC-ESI-MS/MS analysis for O^6^-HE-dG adduct was conducted with the aforementioned Q Exactive HF MS system with the same trapping and analytical columns. The O^6^-HE-dG fraction was loaded to trapping column for 3.75 min before eluting to analytical column for separation. For the detection of O^6^-HE-dG, an inclusion list was comprised of *m*/*z* 312.1303 (O^6^-HE-dG) and *m*/*z* 316.1554 (O^6^-HE-dG-d4) for PRM-MS/MS analysis. LC gradient conditions and MS-PRM parameters are shown in **Supplementary Table 2**.

### Data Analysis

The MS raw data was checked and analyzed with Xcalibur software (Thermo Fisher Scientific). The MS response calibration curves for adduct quantitation were obtained by using the integrated peak area and the known amount ratios of synthetic analytical and internal standards. The quantitative analysis was done by extracting the major fragment ion, as shown in **Supplementary Table 2**, in each corresponding PRM or SRM event using Skyline v24.1.0.199 (MacLean et al. 2010).

### Statistical Analysis

Dose-response regression modeling was performed using GraphPad Prism (version 10.2.0, San Diego, CA. For low-dose exposures (0–1 ppm), a parametric linear regression model was used to characterize the quantitative relationship between EtO exposure and N7-HE-G adduct formation. Mean N7-HE-G adduct levels measured at five defined EtO exposures served as the dependent variable, while EtO exposure concentrations were treated as continuous independent variables. Model selection was guided by visual inspection of residual plots and assessment of lack-of-fit statistics. Regression outputs included slope, intercept, standard errors, and 95% confidence intervals. The model demonstrated a strong goodness-of-fit (R² > 0.98). Assumptions of normality and homoscedasticity were evaluated through residual diagnostics. At higher EtO exposure levels (≥50 ppm), a weighted second-order polynomial regression model was applied to account for the observed nonlinear dose-response relationship in male and female mice. Weighted least squares estimation was used, with weights calculated as the inverse variance of the mean responses, incorporating both sample size and standard error. This approach allowed appropriate modeling of heteroscedasticity across exposure groups.

Pairwise comparisons of N7-HE-G adduct levels were conducted using two-tailed parametric Welch’s t-tests. Specifically, comparisons between each EtO exposure group and the control group were performed to assess dose-related effects. Separately, sex differences (male vs. female) within each exposure group were evaluated using the same statistical approach to account for potential inequality of variances. For each comparison, group means, standard errors, and sample sizes were used to compute t-statistics and corresponding p-values. A two-sided alpha level of 0.05 was applied to determine statistical significance. Pairwise comparisons were conducted for descriptive purposes. No adjustment for multiple comparisons was applied, as pairwise comparisons were considered exploratory.

## RESULTS

### Method Development for the Detection and Quantitation of DNA Adducts

The primary aim of this study is to evaluate the dose response relationships of EtO in the formation of DNA adducts within the mouse tissues. EtO is highly reactive and has a short biological half-life, therefore, its exposure assessment relies on detecting stable biomarkers in hemoglobin and DNA. It is known that EtO can directly react with DNA molecules to form a variety of DNA adducts, including N7-HE-G and O^6^-HE-dG that can be further released for analytical detection after unique experimental preparations (**Scheme 1**). In addition, absolute quantification of DNA adducts can be achieved by monitoring unique fragment ion and utilizing stable isotope-labeled internal standards (Lu et al. 2022). To reveal the molecular dosimetry of DNA adducts in mice exposed to EtO, we conducted a comprehensive experiment to quantify N7-HE-G and O^6^-HE-dG adducts induced directly by EtO exposure (**Figure 1**). The nano-LC–MS/MS method achieved superb sensitivity, with on-column limits of detection of 30 amol and 3 amol for N7-HE-G and O^6^-HE-dG, respectively.

**Scheme 1.**
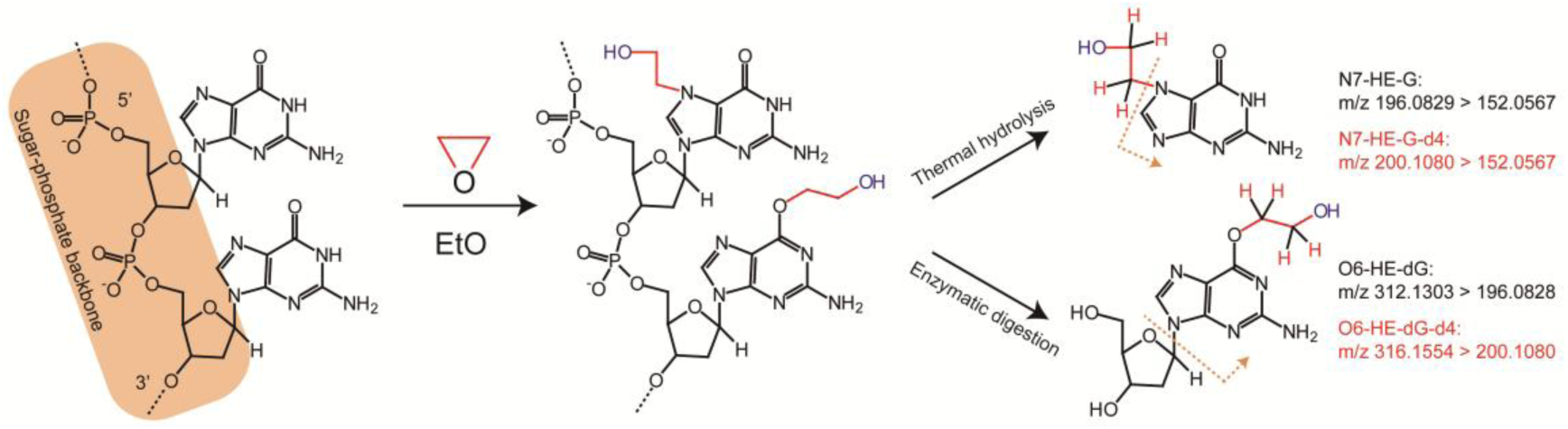
Ethylene oxide (EtO) directly reacts with DNA to form two key DNA adducts. After unique cleavages by thermal hydrolysis and enzymatic digestion, N7-HE-G and O^6^-HE-dG are released for quantitative analysis in mass spectrometry by monitoring corresponding unique fragment ions. The red H indicates stable isotope (deuterium)-labeled internal standards, N7-HE-G-d4 and O^6^-HE-dG-d4, used in this study for accurate absolute quantification.

**Figure 1.**
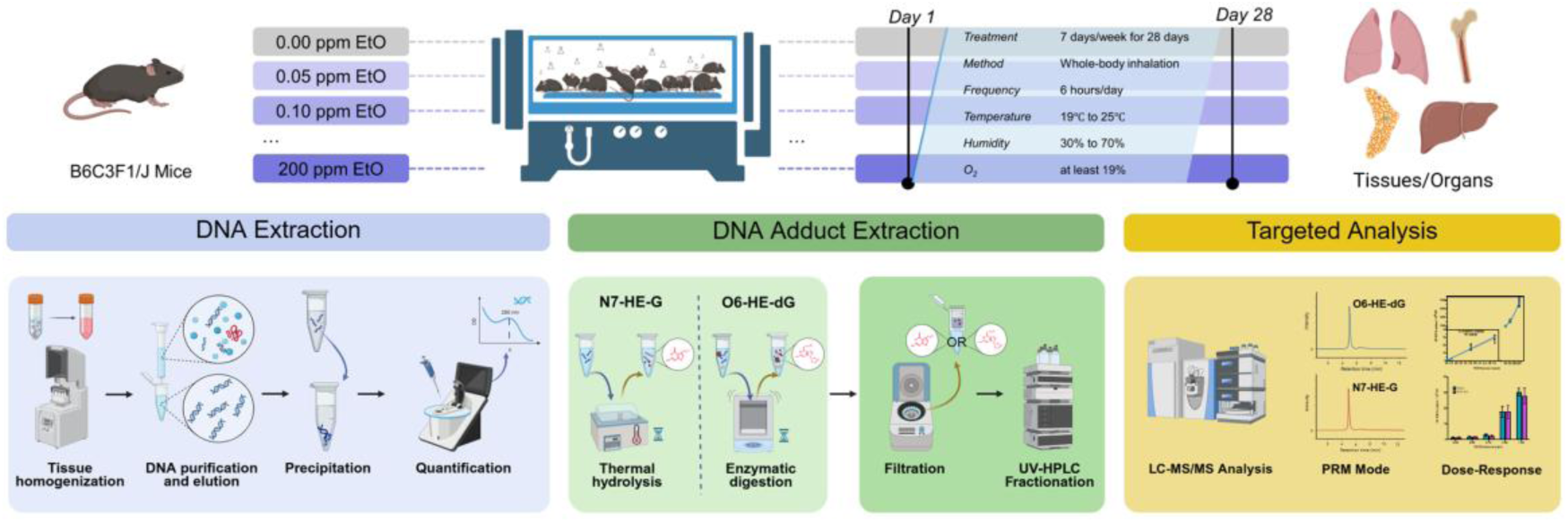
Experimental design and sample processing workflow. B6C3F1/J mice were exposed to different doses and air control via whole body inhalation for 28 consecutive days. Lung, liver, bone marrow and mammary gland tissue were collected after exposure for DNA extraction. Extracted DNA samples were further treated to release N7-HE-G and O^6^-HE-dG adducts by neutral thermal hydrolysis and enzymatic digestion, respectively. Two purification steps including 3 kDa MWCO filtration and HPLC fractionation were conducted to minimize the matrix effect from the complex digested samples before targeted LC-MS/MS analysis of DNA adducts.

### EtO-induced N7-HE-G Levels in Tissues

N7-HE-G adducts in lung, the primary tissue in direct contact with inhaled EtO,demonstrates the representative LC-MS/MS extracted ion chromatograms (XIC) of N7-HE-G and spiked internal standard N7-HE-G-d4 in the lung tissues from male mice exposed to air control, 1 ppm and 200 ppm of EtO (**Figure 2**). Endogenous N7-HE-G adducts were clearly detected in control mice based on the exact mass of expected fragment ion and identical retention time by comparing to its internal standard. The peak area ratio of N7-HE-G over N7-HE-G-d4 was 0.037 from one of those G1 lung samples (**Figure 2A**). A significant increase in N7-HE-G generation was observed with increasing EtO exposure doses. For example, the peak area ratios were 0.836 and 199.439 measured from the mice lung tissues after exposure with 1 ppm (G5) and 200 ppm (G8) of EtO exposure, respectively (**Figure 2B** and **2C**).

**Figure 2.**
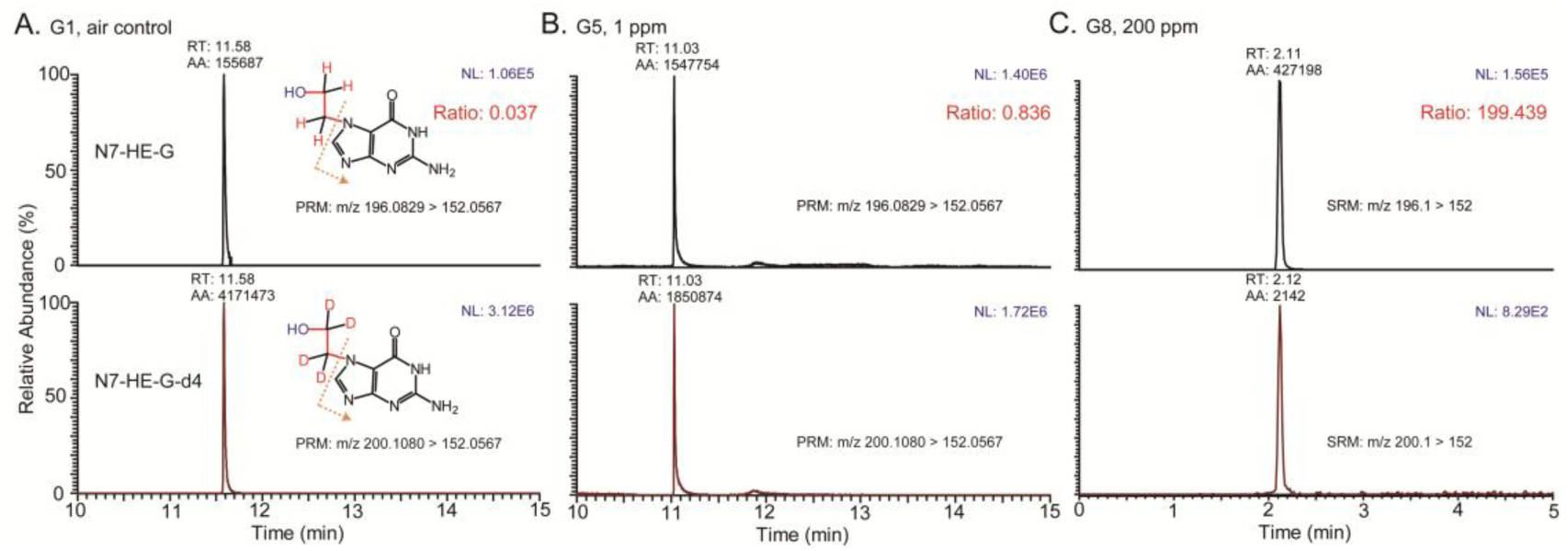
Representative LC-MS/MS PRM (A, B) and SRM (C) extracted ion chromatograms of N7-HE-G (upper panel) and spiked internal standard N7-HE-G-d4 (lower panel) in lungs of male mice exposed to air control (G1), 1 ppm (G5) and 200 ppm (G8) of EtO. Chemical structures of N7-HE-G and N7-HE-G-d4 and their quantifying transition are annotated. The dashed line shows the major fragment ion generated during MS/MS fragmentation for targeted quantification. RT, AA and NL indicate retention time, peak area, and intensity, respectively.

To quantify EtO-induced DNA adducts, the number of N7-HE-G adducts was determined using the peak area ratio of N7-HE-G over its internal standard, N7-HE-G-d4 and further normalized to the amount of DNA subjected to hydrolysis. To more clearly visualize the dose-dependent increase of N7-HE-G, two bar graphs were created using quantified data from male and female mice lung tissues, separating the low (≤1 ppm) and high (≥50 ppm) EtO exposure groups due to the marked difference in N7-HE-G induction between these ranges (**Figure 3**). The quantitative profiles of N7-HE-G adducts in lungs of male and female mice exposed to EtO illustrated a clear dose-disproportionate dose-dependent response in higher EtO concentration exposures (≥50 ppm). Both male and female mice displayed similar increase trends. Moreover, even at the lowest tested dose of 0.05 ppm EtO, a statistically significant increase in N7-HE-G adduct number relative to the air control was detected. The quantified N7-HE-G adducts in lung are further summarized in **Table 1**. These data demonstrate that N7-HE-G adducts were generally proportional to external EtO exposure up to 100 ppm EtO, and increased dose-disproportionately between 100 and 200 ppm EtO.

**Figure 3.**
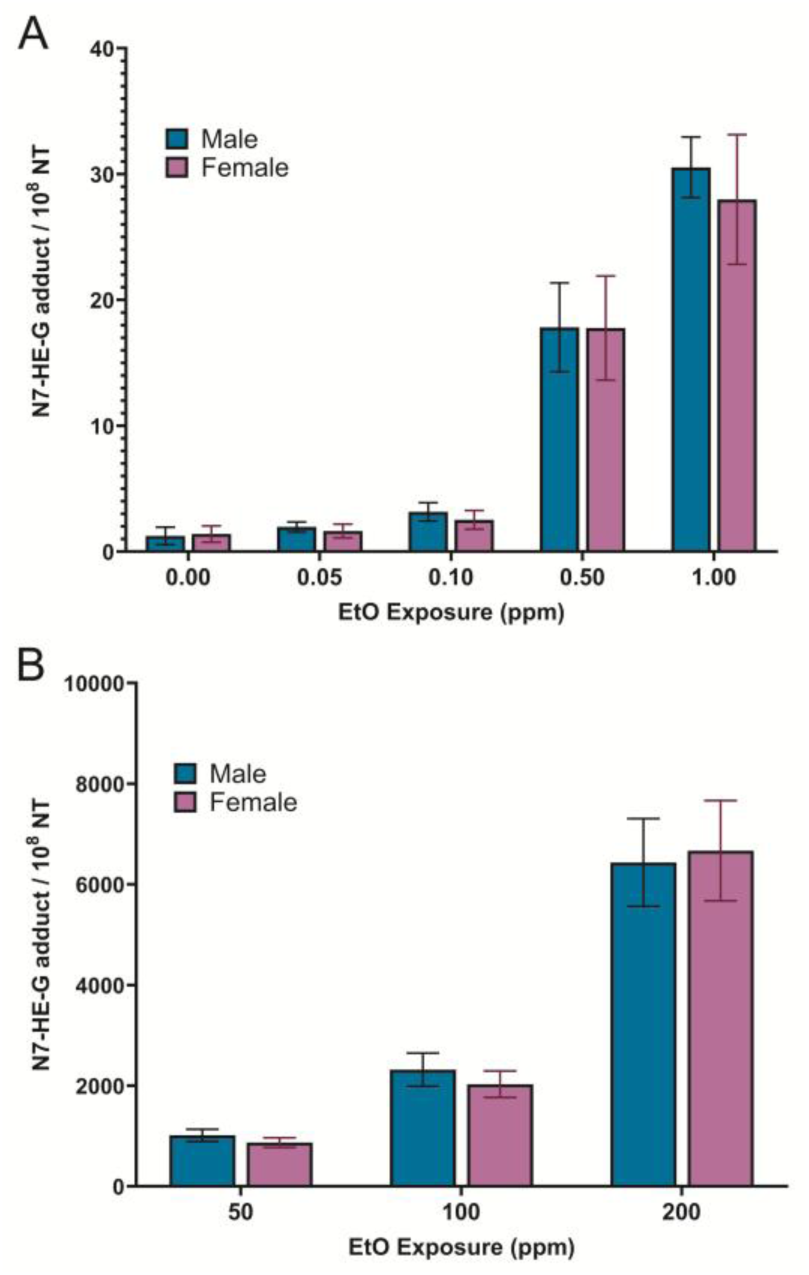
N7-HE-G adducts (per 10^8^ NT, nucleotides) in lungs of mice exposed to low (≤1 ppm, A) and high (≥50 ppm, B) EtO exposures. Each data point represents the mean ± standard deviation (SD) at a given exposure level.

**Table 1.**
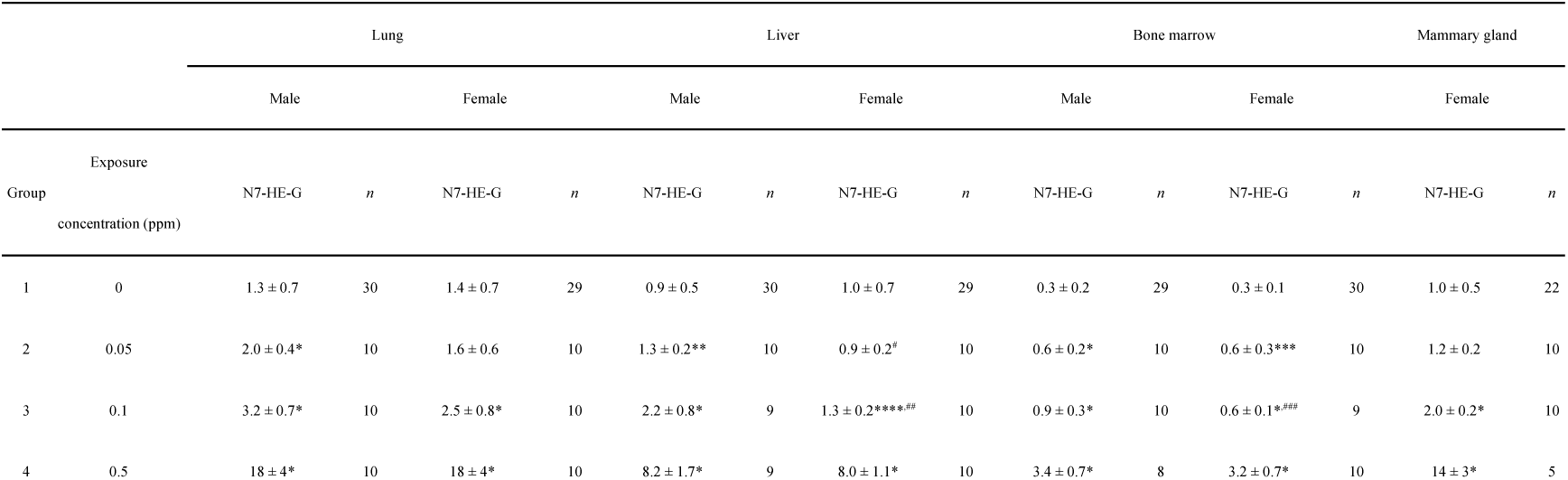
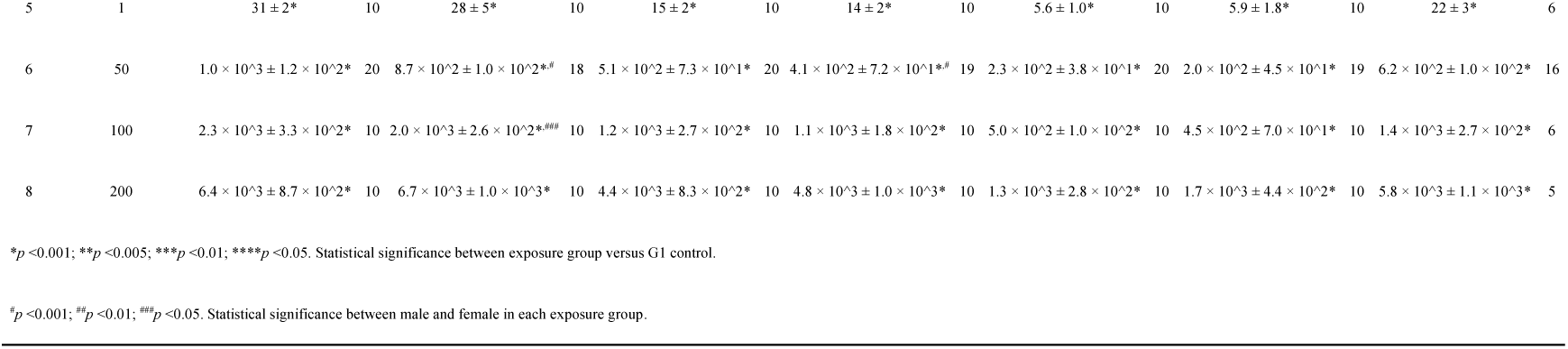
N7-HE-G adduct numbers (10^8 NT) quantified in the lung, liver, bone marrow, and mammary gland collected from mice exposed to EtO. Values are presented as mean ± SD and rounded to an appropriate level of precision. Scientific notation is used where appropriate.

The dose-response plots illustrating the relationship between EtO exposures and N7-HE-G adducts in male and female lung is further depicted in **Figure 4**. A biphasic dose-response relationship in N7-HE-G formation was clearly observed in both sexes following EtO exposure. At low doses (0–1 ppm), a clear linear relationship was presented, as shown in the inset linear regression plots (R² = 0.993 for males; R² = 0.981 for females). At a higher concentration range (50–200 ppm), an increasingly steeper non-linear-dose-response pattern was observed. These results demonstrated a strong positive correlation between EtO exposure and N7-HE-G formation in mouse lung.

**Figure 4.**
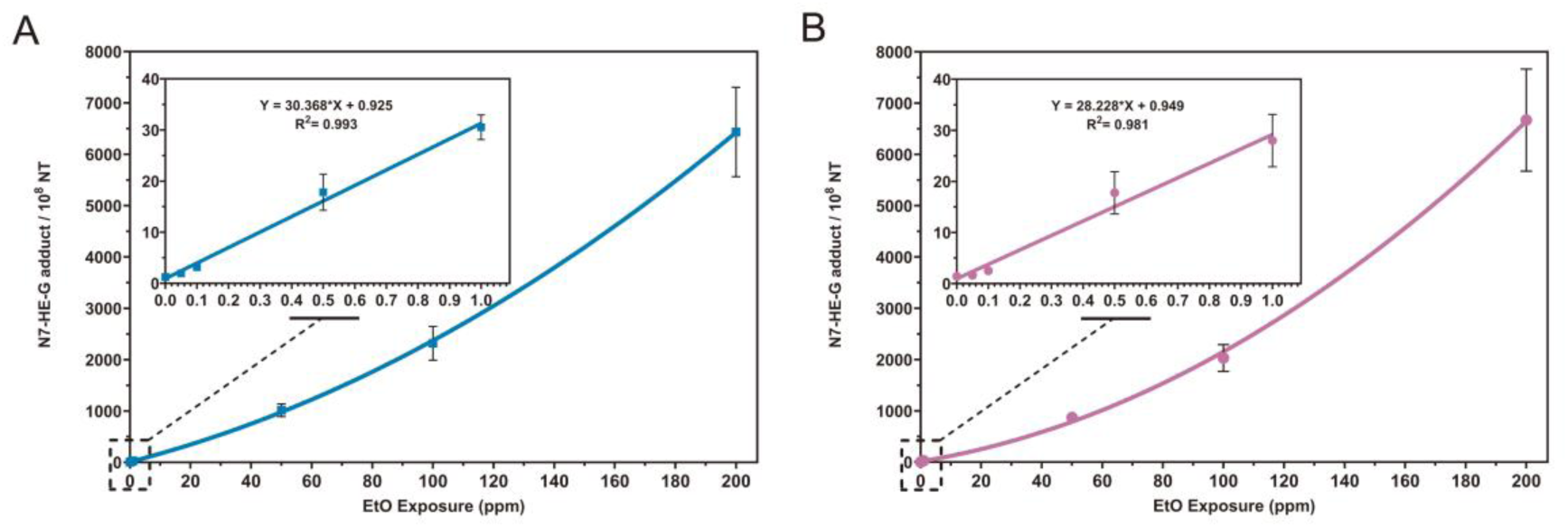
Dose-response curves of N7-HE-G adducts in lungs of male (A) and female mice (B) across the full exposure range (0–200 ppm). N7-HE-G adducts (per 10^8 NT, nucleotides) were quantified in lung tissue samples of male and female mice after exposure to various concentrations of EtO. Each data point represents the mean ± standard deviation (SD) at a given exposure. Insets show the linear regression fit for the low-dose range (≤1 ppm), with corresponding equations and R² values.

N7-HE-G adducts were also analysed in liver, bone marrow and mammary gland. This analysis aimed to evaluate whether EtO exposure leads to systemic distribution in DNA adduct accumulation beyond the primary site of contact. The quantitative results of N7-HE-G adduct from those examined tissues are also summarized in **Table 1**. Notably, all tissues exhibited the presence of adducts in the control non-exposed mice, an observation consistent with endogenous production associated with normal body metabolism (Kirman et al. 2021; Kirman et al. 2025). Bone marrow had the lowest endogenous N7-HE-G adduct level (0.319 and 0.305/10^8 NT for male and female mice, respectively) among the four examined organs. The order of N7-HE-G adduct from highest to lowest among tissues was lung, mammary gland, liver and bone marrow, especially in the higher dose groups (≥0.5 ppm). The dose-response curves of N7-HE-G adducts presented similar patterns between tissues and male/female sex, i.e. a linear increase in the low-dose range with a shallower slope (≤1 ppm) versus sublinear increase with a steeper slope in the high-dose range (≥50 ppm). Moreover, statistical analysis was performed to determine whether significant differences existed between each EtO exposure group and the air control, and to examine potential sex-related effects (**Table 1** and **2**). Overall, there is a significant difference between each EtO exposure group and the air control, except 0.05 ppm samples from lung, liver and mammary gland in female mice. Sex-dependent differences in adduct levels were minimal and not statistically significant in most tissues and exposure groups.

### O^6^-HE-dG Levels in Mice Exposed to High Dose of EtO

**Figure 5** displays the representative LC-MS/MS XIC of the promutagenic O^6^-HE-dG and spiked internal standard O^6^-HE-dG-d4 measured in lung from male mice exposed to air control (G1) and 200 ppm of EtO (G8). Notably, endogenous O^6^-HE-dG adducts, using up to 20 μg DNA for hydrolysis and our sample preparation methodology, were not detected in any tissues from air control mice (**Figure 5A and Table 2**). In addition, O^6^-HE-dG adducts were consistently not detected in any samples from G2 (0.05 ppm) to G5 (1 ppm) exposure groups, indicating that formation of this promutagenic adduct requires relatively high levels of EtO exposure.

**Figure 5.**
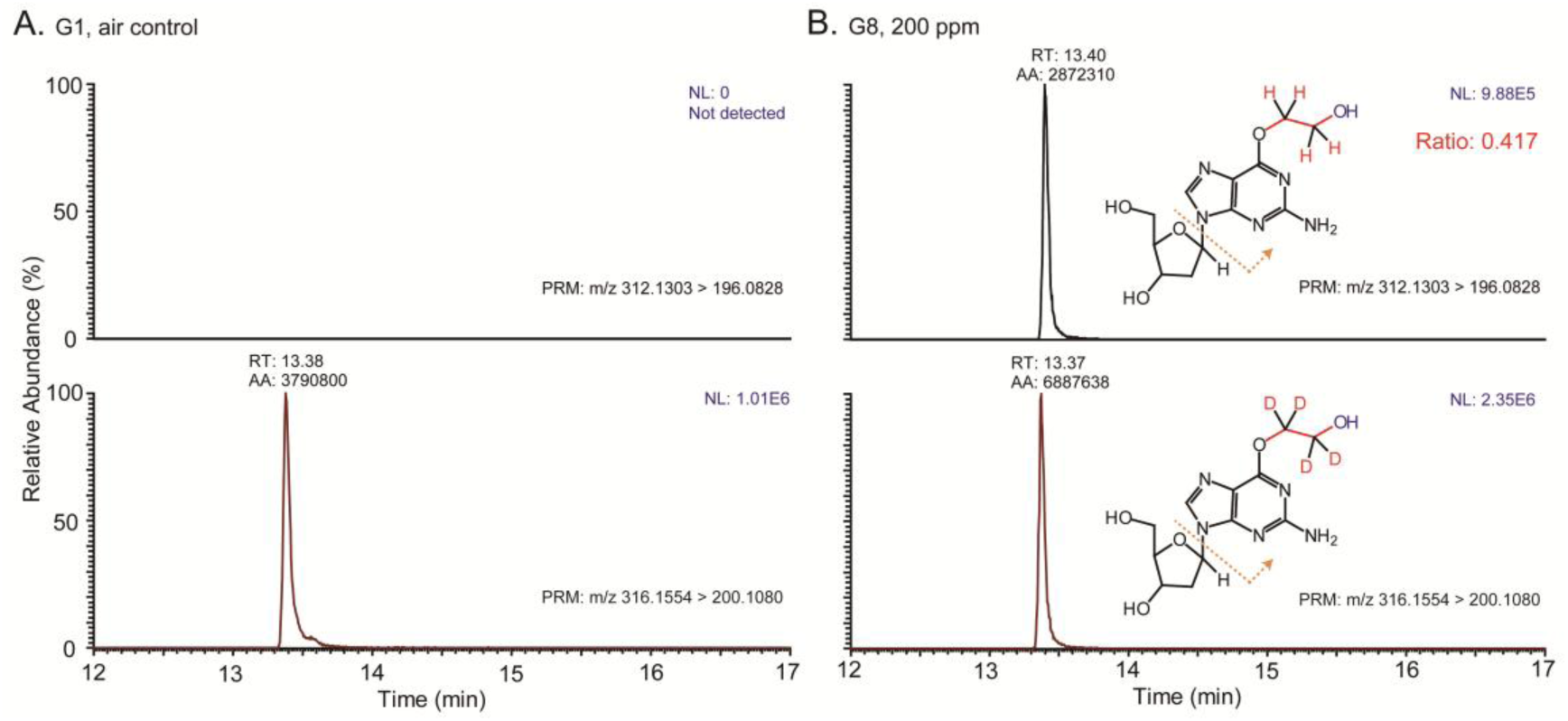
Representative LC-MS/MS PRM extracted ion chromatograms of O^6^-HE-dG (upper panel) and spiked internal standard O^6^-HE-dG-d4 (lower panel) in lungs of male mice exposed to air control (G1) and 200 ppm (G8) of EtO. Chemical structures of O^6^-HE-dG and O^6^-HE-dG-d4 and their quantifying transition are annotated.

**Table 2.**
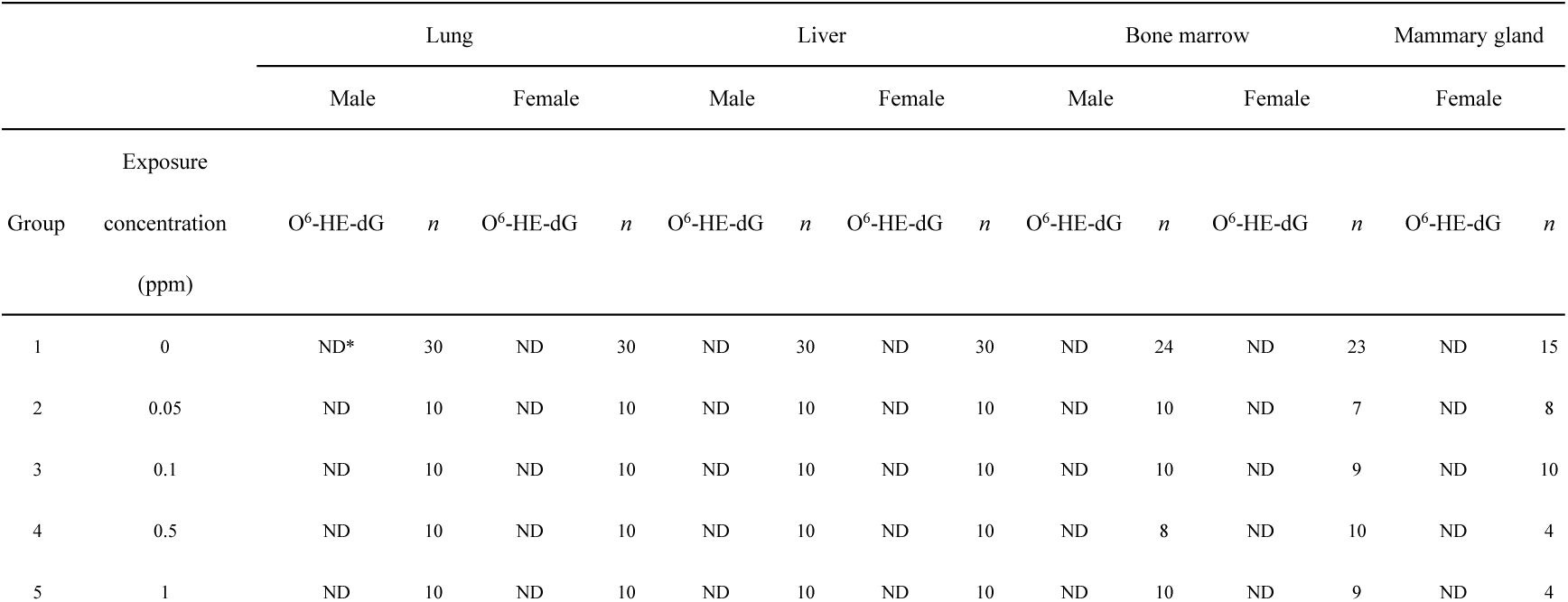
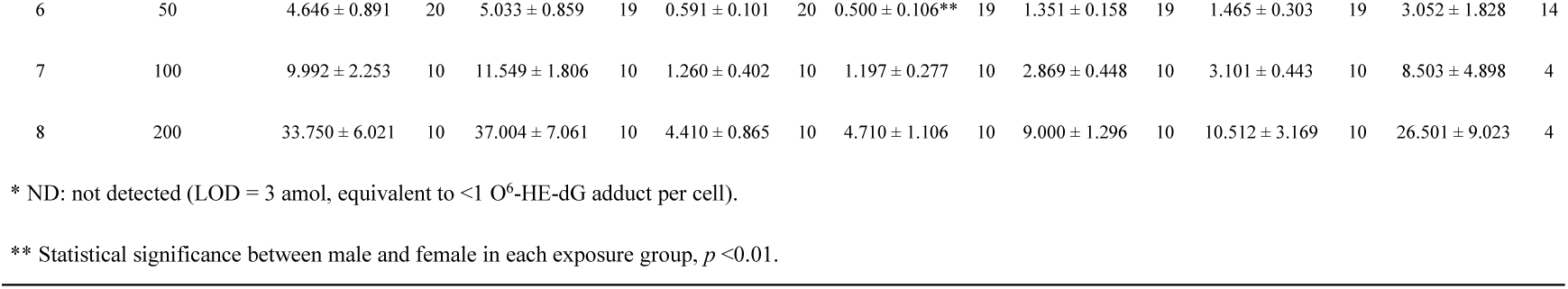
O^6^-HE-dG adduct numbers (10^8 dG) quantified in the lung, liver, bone marrow, and mammary gland collected from mice exposed to EtO.

In lung, O^6^-HE-dG adducts became quantifiable starting at 50 ppm of EtO exposure, with mean levels of 4.646 (male) and 5.033 (female) per 10^8 dG and increased dose-disproportionately at 200 ppm (33.750 and 37.004 per 10^8 dG). Notably, the analytical sensitivity of our current method for O^6^-HE-dG detection is 3 amol, which is equivalent to less than one adduct per cell. Lung exhibited the highest O^6^-HE-dG formation, followed by mammary gland, bone marrow, and liver. **Figure 6** shows dose-response curve of O^6^-HE-dG adducts in lungs of male and female mice following EtO exposure. It is consistent with the dose-response pattern of N7-HE-G in high-dose range (≥50 ppm), showing a sublinear increasing trend. Similarly, this apparent threshold and increasing dose-response pattern was also exhibited in the other examined tissues (**Table 2**). These results collectively indicate that O^6^-HE-dG formation is restricted to mice exposed only to high doses of EtO.

**Figure 6.**
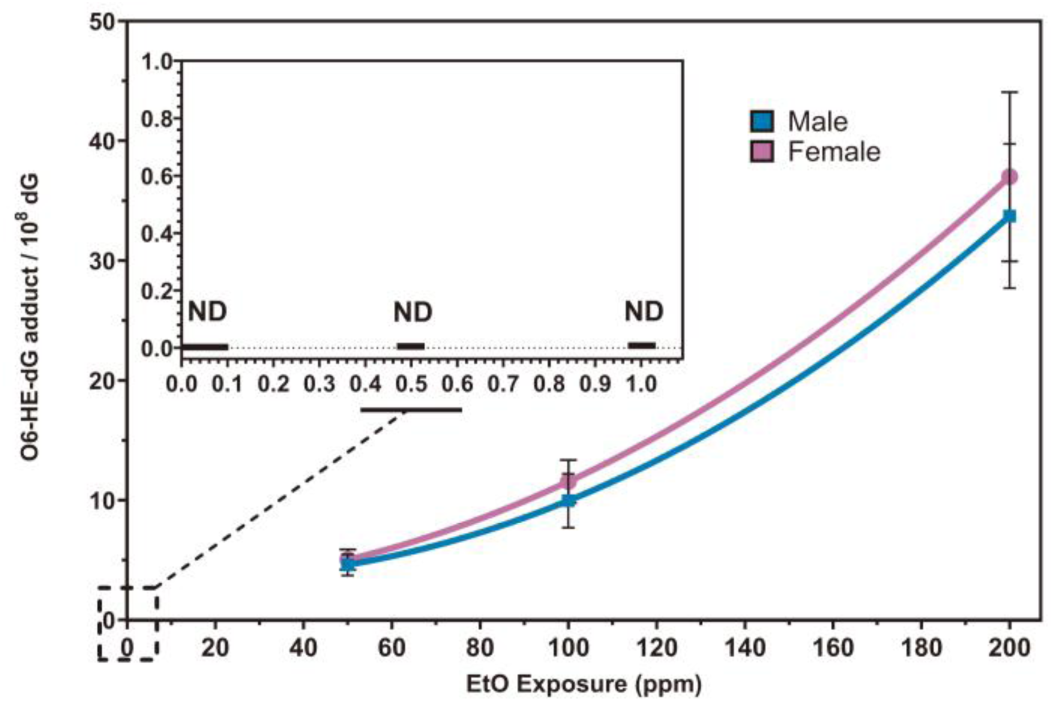
Dose-response curves of O^6^-HE-dG adduct formation in lungs of male and female mice across the full exposure range (0–200 ppm). Each data point represents the mean ± standard deviation (SD) at a given exposure level. O^6^-HE-dG adducts were not detected (ND) in the low-dose range (≤1 ppm).

Previous analyses have reported that N7-HE-G is formed at approximately 300-fold higher levels than O^6^-HE-dG in rats exposed to 300 ppm for 1-4 weeks (Walker et al. 1992). Using methodology with substantively improved sensitivity for measurement of O^6^-HE-dG and improved titration of the exposure at which O^6^-HE-dG is first detected (50 ppm), the ratios of N7-HE-G to O^6^-HE-dG formation for each tissue and respective male and female gender at the 50 ppm exposure (**Tables 1** and **2**) are lung: 220, 173; liver: 866, 816; bone marrow: 170,140; and female mammary gland: 202. The higher ratio of N7-HE-G to O^6^-HE-dG in the liver is likely a reflection of efficient repair of O^6^-HE-dG adduct by methylguanine methyltransferase (MGMT) enzyme in this tissue. These adduct ratios are in general agreement with those previously reported and vary within an order magnitude across the examined tissues.

### Strong Correlations between Tissue DNA and Blood Protein Adducts

The overall protocol of the current study also included quantification of systemic EtO exposures as measured by formation of EtO-induced N-(2-hydroxyethyl)-L-valine (HE-V) hemoglobin adducts in blood collected from the same exposed mice used in this study (Liu et al. submitted). Interestingly, the HE-V dose-response pattern presents high similarity with those of various tissue N7-HE-G patterns across all EtO exposure groups. To investigate whether blood protein adducts could serve as surrogates for tissue-specific DNA damage, we assessed the statistical correlation between hemoglobin HE-V and N7-HE-G in tissues. A strong and positive linear correlation was observed between blood HE-V and N7-HE-G adduct levels in lung, liver, bone marrow and mammary gland (p < 0.001, **Figure 7**).

**Figure 7.**
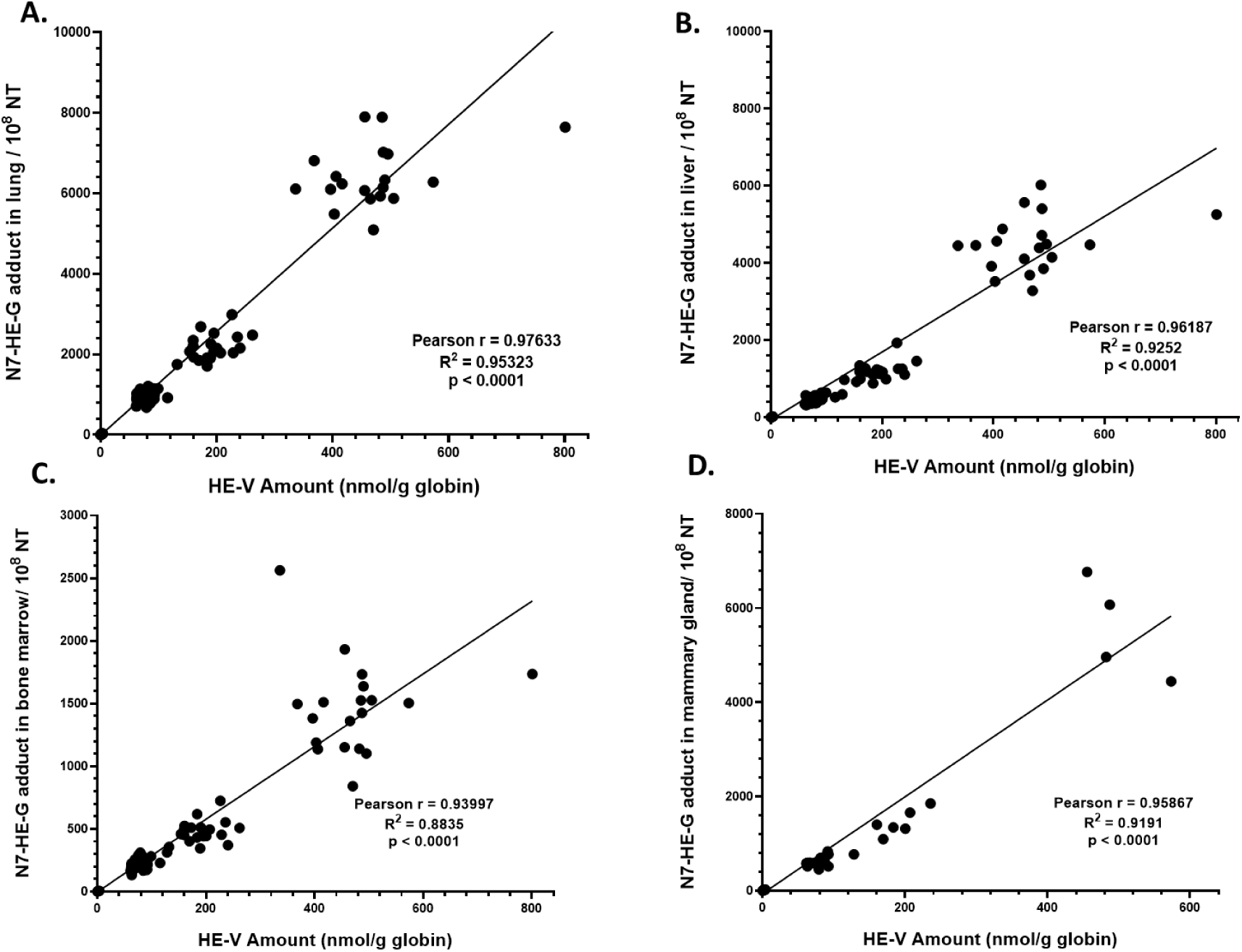
Correlation between blood HE-V levels and DNA adducts in EtO-exposed mice (A: lung; B: liver; C: bone marrow; and D: mammary gland). Pearson correlation between N7-HE-G adducts in lung and HE-V levels in blood across all EtO exposure groups. HE-V data collected from same experimental tissues (Liu et al. submitted).

## DISCUSSION

A key objective of the current study was to characterize the dose-response patterns of EtO-induced DNA adduct formation, a molecular initiating MOA event hypothesized for EtO carcinogenesis, in order to provide experimental dose-response data guiding selection of a human cancer risk assessment model whose dose-response shape is biologically plausible when considered against EtO’s hypothesized MoA. In order to achieve this goal, sensitive methods were developed to quantify DNA adducts in bone marrow, mammary tissue, lung and liver. This study demonstrates that the shape of the non-mutagenic N7-HE-G adduct dose-response is linear at EtO exposures ≤1 ppm, while pro-mutagenic O^6^-HE-dG adducts were detected starting only at 50 ppm and greater exposures. At EtO exposures ≥50 ppm, the dose-response slopes for both adducts increased disproportionately with increasing EtO exposures. Importantly, in addition to being the first report of describing N7-HE-G adducts at and below 1 ppm EtO, the absence of detectable O^6^-HE-dG adducts at <1 ppm EtO is not attributable to insufficient analytical sensitivity. To our knowledge, the current method provides the lowest reported limit of detection for O^6^-HE-dG (3 amol), exceeding the sensitivity of previously published LC-MS/MS assays, including those reported by Zhang et al (Zhang et al. 2015). Previously reported methods achieved limits of quantitation in the low-femtomole per milliliter range, whereas the present method achieves attomole-level sensitivity. Assuming approximately 6 pg of genomic DNA per cell, this detection limit corresponds to fewer than one O^6^-HE-dG adduct per cell.

While N7-HE-G formation is highly responsive and abundant, its biological significance as a mutagenic lesion is limited. The N7 position does not directly participate in Watson-Crick base pairing, and this type of adduct is prone to spontaneous depurination, potentially leading to apurinic/apyrimidinic (AP) sites rather than base mispairing (Swenberg et al. 2011). Thus, although N7-HE-G provides an excellent dosimetric marker similar to HE-V in blood, it is unlikely to be a major contributor to EtO-induced mutagenesis or carcinogenesis.

In contrast, O^6^-HE-dG was only detected at higher EtO exposures (≥50 ppm) (**Figure 6 and Table 2**). Although far less abundant than N7-HE-G, O^6^-HE-dG is highly promutagenic due to its mispairing potential with thymine during DNA replication, resulting in G:C ➔ A:T transition mutations (Delaney and Essigmann 2001; Mazon et al. 2010). The absence of O^6^-HE-dG at low-dose exposures (0.05–1 ppm), despite readily detectable N7-HE-G formation and the exquisite sensitivity of our technique, underscores the disparity between internal dose measured as HE-V or N7-HE-G and effective genotoxic dose.

The two most recent cancer risk assessments for EtO are based on the same epidemiologic study of sterilant workers conducted by the National Institute of Occupational Safety and Health (NIOSH) but result in cancer risk estimates with three orders of magnitude difference (TCEQ 2020; USEPA 2016; Valdez-Flores et al. 2025). Both TCEQ and USEPA assume a linear low-exposure extrapolation based on a presumed mutagenic mode of action but apply very different dose-response models to the NIOSH data. Both agencies applied Cox Proportional Hazards (CPH) models to the NIOSH data. However, TCEQ and USEPA applied different forms of CPH model that had comparable statistical significance. EPA selected a 2-piece linear spline model exhibiting an extremely steep linear slope at low exposures that shallows at higher exposures. In contrast, TCEQ relied on a shallower single-slope log-linear dose-response model that is nearly linear across all exposures in the NIOSH study (Kirman et al. 2025; TCEQ 2020; USEPA 2016; Valdez-Flores et al. 2025).

Although it can be argued based on statistical principles alone that the TCEQ model approach is more parsimonious (simpler model), consideration of biological plausibility should be the primary basis for selecting statistical models, especially when both models have similar fit to the observed individual data. The EPA Science Advisory Board’s review of the EPA IRIS assessment emphasized that “any model that is to be considered reasonable for risk assessment must have a dose response form that is both biologically plausible and consistent with the observed data.” (USEPA SAB 2015).

EPA carcinogen risk assessment guidelines states that “[i]f dose-response analysis of nontumor key events is more informative about the carcinogenic process for an agent, it can be used in lieu of, **<u>or in conjunction with</u>**, tumor incidence analysis for the overall dose-response assessment. (emphasis added)” (USEPA 2005). Thus, examination of the dose response relationships of DNA adduct formation in the current study offers a biologically plausible and MOA-informed insights into which of the epidemiology-based statistical dose-response models reliably predicts low-exposure EtO cancer risks. In this same vein, a National Academy of Science review of the TCEQ cancer risk approach concluded that MOA data focused on identifying the comparative *in vivo* dose-response relationships of EtO DNA adducts to those of apical genotoxicity would be valuable in guiding the selection of a statistical dose-response model that most appropriately reflects a biologically plausible estimation of low-exposure EtO cancer risks (National Academies of Sciences 2025).

Overall, the DNA adduct dose-response data are completely inconsistent with the postulated EtO dose-response shape derived by EPA from statistical modeling of empirical epidemiology cancer data, i.e., a very steep initial increase in risk in the low EtO exposure range that shallows at higher occupational exposures (USEPA 2016). In contrast, the EtO DNA adduct dose-responses support a biologically-plausible informed selection of a shallower single-slope linear cancer dose-response model across the entire range of low to high EtO exposures (TCEQ 2020; Valdez-Flores et al. 2025). The implications of such dose-response model choices to EtO cancer risk assessment are dramatically illustrated by the three order of magnitude lower cancer risk predicted by the TCEQ (TCEQ 2020), whose cancer dose-response model assumed an essentially single-slope linear dose-response based on the standard log-linear Cox Proportional Hazards modeling of the same EtO occupational cancer dataset used by the EPA.

Another key objective of the current experiment protocol was to inform the shape of the EtO dose response inclusive of three levels of increasing biological MOA information: 1) systemic EtO exposure measured by blood HE-V (Liu et al. submitted); 2) DNA adducts (this study); and 3) apical manifestation of cytogenetic (MN assay) and mutagenic activity (*Pig-a*) in blood reticulocytes and mature erythrocytes (Gollapudi 2023). The shape of the dose-response for plasma HE-V exactly paralleled that of cancer target organ N7-HE-G adducts (i.e., linear at ≤1 ppm, upward-bending at ≥50 ppm), while apical genotoxicity was largely apparent only at 200 ppm EtO. These integrated dose-response responses are consistent only with a conservatively-assumed shallow single-dose linear dose response at all three levels of biological MOA considerations.

The EtO DNA adduct and genotoxicity findings associated with this study protocol are consistent with similar higher-dose EtO induced genotoxicity dose-response patterns reported in the literature, all of which lead to a conclusion that EtO is a weakly active genotoxicant (Gollapudi et al. 2020; Hartwig et al. 2020; Pottenger et al. 2019). Detection of the promutagenic O^6^-HE-dG exclusively in the higher-dose groups aligns with a hypothesis that EtO may exhibit negligible mutagenic activity attributed in part to efficient low-exposure repair by O^6^-alkylguanine-DNA alkyltransferase (AGT), which is a mechanism that is saturated in high exposure scenarios (Jenkins et al. 2005). These results indicate that the shallow nearly linear model selected by TCEQ (TCEQ 2020), while still representing a conservative model choice for EtO cancer risk assessment, has greater biological plausibility than the 2-slope linear model with initial very steep slope.

Interestingly, the mammary gland, despite being one of the most distal tissues from the site of inhalation, exhibited comparable N7-HE-G adduct levels to the lung at 200 ppm EtO exposure, and ranked second among the four analyzed tissues at most other exposure levels. A similar pattern was also observed for O^6^-HE-dG, with the mammary gland consistently exhibiting the second highest adduct levels across all high-dose exposure groups. Although these data suggest alignment with existing animal and epidemiological evidence, such an interpretation is tempered by the observation that the relatively crude punch-biopsy collection of mammary tissue likely resulted in admixing with unknown quantities of surrounding adipose tissue.

The dose-response patterns for HE-V, N7- and O^6^-DNA adducts and genotoxicity are also aligned with PBPK-modeled EtO toxicokinetic dose-response patterns in mice, rats and humans (Fennell and Brown 2001). Importantly, PBPK modeling predicted EtO blood toxicokinetics were linear and with similar blood concentrations observed for all species up to a 100-ppm exposure. However, and consistent with the EtO HE-V data, mouse blood concentrations increased dose-disproportionately between 100 and 200 ppm, while rat and human concentrations exhibited continuing linear behavior at exposures ≥300 ppm. The higher-exposure upward bend in mouse toxicokinetics was attributed to preferential depletion of glutathione in mice resulting in reduced detoxification capacity mediated by EtO glutathione conjugation (Filser and Klein 2018). Thus, as with the HE-V, DNA adduct and genotoxicity data, systemically-modeled PBPK EtO blood concentration data offer further evidence that EtO conservatively operates by single-slope linear dose responses across a wide range of EtO exposures in multiple mammalian species.

The strong correlation observed between blood HE-V protein adducts and tissue N7-HE-G adducts underscores the utility of HE-V as a minimally invasive biomarker of internal EtO exposure. HE-V is an effective biomarker for EtO biomonitoring due to its stability, detectability, and continuously linear dose response between low ppb environmental to high ppm occupational exposures (Kirman et al. 2025). Our findings clearly extend its application by demonstrating that HE-V levels not only reflect total EtO burden but also correlate with N7-HE-G exposure in the tissues. This is especially relevant for retrospective exposure assessment in epidemiological studies, where access to tissue samples is limited and blood biomarker, HE-V, provides a practical alternative. To the best of our knowledge, we showed, for the first time, significantly increased N7-HE-G adducts in the low exposure range (≤1 ppm). Regression analysis confirmed a strong linear correlation between N7-HE-G formation and low-dose EtO exposure (≤1 ppm), and adducts were sharply increased in the high-dose exposure (**Figure 4**). This specific pattern was observed in all analyzed tissues from both male and female mice. Moreover, our data also revealed tissue-specific variations in adducts affected by EtO exposure. Following inhalation exposure, EtO is readily absorbed through the pulmonary alveoli and enters the systemic circulation, which facilitates delivery of reactive EtO molecules to distalf liver, bone marrow, and mammary gland tissues Lung, as a primary site of contact, consistently showed higher N7-HE-G and O^6^-HE-dG levels compared to liver and bone marrow. Interestingly, the mammary gland, despite being one of the most distal tissues from the site of inhalation, exhibited comparable N7-HE-G adduct levels to the lung at 200 ppm EtO exposure, and ranked second among the four analyzed tissues at most other exposure levels (**Table 1**). A similar pattern was observed for O^6^-HE-dG, with the mammary gland consistently exhibiting the second highest adduct levels across all high-dose exposure groups. These findings support the biological plausibility that high EtO exposures may contribute to development of both lung and breast cancers, in alignment with existing animal and epidemiological evidence. However, such an interpretation needs to be tempered by the observation that mammary tissue was obtained by a punch biopsy of nipples that was likely admixed with the surrounding adipose tissue, skin and hair.

## CONCLUSION

Using stable isotope-labeled internal standards and highly specific, analytically sensitive LC-MS/MS quantification, our study characterized the dose-response patterns of both non-mutagenic N7-HE-G and pro-mutagenic O^6^-HE-dG DNA adducts across multiple tissues in both sexes of mice. These analyses followed inhalation exposure to a 4000-fold range of EtO concentrations, spanning potential low parts-per-billion environmental exposures up to substantially higher parts-per-million levels typical of occupational settings. The results revealed a linear dose-response pattern for N7-HE-G adduct formation that was highly correlated with the levels of N-(2-hydroxyethyl)-L-valine (HE-V), supporting the use of HE-V as a surrogate biomarker of EtO dose to DNA. In contrast, the dose-response pattern for pro-mutagenic O^6^-HE-dG adducts provided critical molecular-level MOA data, essential for selecting a biologically plausible dose-response model. Taken together, these findings support the conclusion that the cancer risk to the general population from EtO exposure is conservatively modeled by assuming a MOA that is characterized by a shallow single-slope linear dose-response relationship.

## Conflict of interest

The authors CWL, JP, JF, HZ, XW and KL declare they have no actual or potential competing financial interests. Authors BBG, AAL and JS contributions were supported by contracts from the American Chemistry Council to Exponent (AAL, JB) or directly to BBG.

## Acknowledgements

The research was partially supported by the UNC Superfund Research Program (P42ES031007), UNC Center for Environmental Health and Susceptibility grant (P30-ES779 010126), and a gift from the American Chemistry Council. We are grateful to Dr. Joanna Klapacz for her contributions to the conceptualization and design of the study, and to Chris Kirman for his technical review of the manuscript.

## Abbreviations

EtO: ethylene oxide
N7-HE-G: N7-(2-hydroxyethyl)guanine
O^6^-HE-dG: O^6^-(2-hydroxyethyl)-2′-deoxyguanosine
HE-V: N-(2-hydroxyethyl)-L-valine
PRM: parallel reaction monitoring
HCD: higher-energy collisional dissociation
LC-MS: liquid chromatography-mass spectrometry
MS/MS: tandem mass spectrometry
MWCO: molecular weight cut-off
XIC: extracted ion chromatogram

