## Supplemental Method and Tables for "Molecular Dosimetry of DNA Adducts in Mice Exposed to Ethylene Oxide"

The inhalation exposure methodology has been described in detail elsewhere (Liu et al. submitted) and is summarized briefly below.

#### Experimental Animals and Housing

Male and female B6C3F1/J mice (The Jackson Laboratory) were acclimated and assigned to treatment groups (G1–G8) using a stratified randomization scheme. Mice were 9–12 weeks old at the initiation of Phase I and 10–11 weeks old for Phase II. Animals were identified via subcutaneous chips and housed according to the *Guide for the Care and Use of Laboratory Animals*. Environmental conditions were maintained at 20°C–26°C with 30–70% relative humidity and a 12-hour light/dark cycle. Certified Rodent Lab Diet 5002 and municipal tap water (purified via reverse osmosis and UV) were available *ad libitum*, except during exposure periods when a water substitute (HydroGel® Recovery) was provided.

#### Inhalation Exposure System and Regimen

EtO was administered daily via whole-body inhalation (6 h/day) for 28 consecutive days in 1000-L stainless-steel and glass chambers. Exposures were conducted in two phases to address technical challenges in Phase I regarding steady-state concentration generation for the 0.05 and 0.1 ppm groups. Consequently, Phase II utilized a serial chamber design (incorporating a 500-L sampling chamber) to mitigate interferences from animal dander and humidity. Target and measured chamber concentrations for each phase and sex are detailed in **Table S1**, demonstrating high precision in Phase II (e.g., 0.050 ± 0.0027 ppm for Phase II G2 males). Dynamic chamber conditions were maintained with ≥10 air changes per hour and an oxygen content of 20.9%.

**Table S1**. Target and Experimental Concentrations of EtO in Animal Exposures

| Target Exposure Conc. (ppm) | Phase | Male Mean Exposure Concentration (Stdev) | *n* (male) | Female Mean Exposure Concentration (Stdev) | *n* (female) |
|---|---|---|---|---|---|
| 0.00 | 1, 2 | 0 (0.0) | 30 | 0 (0.0) | 30 |
| 0.05 | 2 | 0.050 (0.0027) | 10 | 0.050 (0.0029) | 10 |
| 0.10 | 2 | 0.104 (0.0064) | 10 | 0.103 (0.0061) | 10 |
| 0.50 | 1 | 0.531 (0.1279) | 10 | 0.512 (0.1281) | 10 |
| 1.00 | 1 | 0.961 (0.3238) | 10 | 0.918 (0.1723) | 10 |
| 50 | 1,2 | 51 (5.5) | 20 | 50 (2.6) | 19 |
| 100 | 1 | 98 (8.5) | 10 | 97(6.0) | 10 |
| 200 | 1 | 202 (17.4) | 10 | 200 (16.3) | 10 |

**Tissue Collection and Preservation**

Within two hours of the final exposure, mice were euthanized rotating across dose groups to ensure temporal consistency. Samples of lung (right lung), liver (left lateral lobe), blood, bone marrow, and mammary gland were collected. Bone marrow was harvested from both femurs into fetal bovine serum (FBS), centrifuged at 2300 rpm (4°C, 15 min), and the cell pellet was preserved. Mammary tissue was collected using a 5 mm biopsy punch from the posterior four nipples; hair and skin were removed prior to freezing. All tissues were flash-frozen in liquid nitrogen and stored at –70°C before overnight shipment to the University of North Carolina for DNA and protein adduct analysis.

**Evaluation of Inter-batch Reproducibility and Data Integration**

Due to technical issues in generating steady-state concentrations of 0.05 ppm (G2) and 0.1 ppm (G3) EtO during the initial phase (Phase I), a second phase (Phase II) was performed, including only G1, G2, G3, and G6 to ensure exposure accuracy. While DNA adducts were quantified for all animals, G2 and G3 data from Phase I were excluded from the final dose-response analysis between EtO exposure and DNA adduct formation. To ensure the statistical robustness of the combined dataset, we conducted a rigorous comparative analysis of DNA adduct levels between experimental Phase I and Phase II using the common exposure cohorts (0 and 50 ppm EtO), as shown in **Figure S1** and **S2**. In the 0 ppm control groups, some observed statistical variations in baseline N7-HE-G levels primarily reflect the high sensitivity of our analytical platform in detecting subtle, endogenous biological fluctuations common at near-baseline concentrations. These minor shifts in baseline values are anticipated in large-scale inhalation studies and do not impact the interpretability of the overall dose-response relationship.

The reliability of the integrated data is further evidenced by the high degree of reproducibility observed at the 50 ppm exposure level. For instance, N7-HE-G concentrations in the male bone marrow and liver showed exceptional stability between phases ($p = 0.853$ for both tissues). Similarly, the $O^6$-HE-dG adduct, a critical marker for mutagenic risk, demonstrated remarkable consistency across all evaluated tissues ($p > 0.13$), confirming the high precision of the analytical methodology across independent experimental runs. Most importantly, the dose-dependent kinetic trends remained highly parallel across both phases, with EtO-induced adduct levels at 50 ppm consistently exceeding baseline values by several orders of magnitude.

The integration of G1 and G6 datasets across both phases provides an exceptionally robust statistical foundation for the curve's baseline and high-dose anchor points. This enhanced statistical power, combined with the demonstrated inter-batch reproducibility, ensures a more stable and definitive characterization of the dose-response relationship, particularly when interpolating the biological effects within the critical low-dose region. Furthermore, the robust linearity observed in the low-dose region (0–1 ppm, $R^2$ >0.98, **Figure 4**), supported by these integrated large-sample anchor points, underscores the high precision of our dose-response modeling. This alignment confirms that the biological response to EtO is highly predictable across multiple orders of magnitude, justifying the use of combined anchor points to definitively characterize the critical low-dose landscape. Taken together, the high reproducibility of adduct formation at effective dose levels and the consistency of the overall kinetic response support the integration of Phase I and Phase II datasets. This combined approach maximizes the statistical power necessary for a definitive characterization of the dose-response relationship in the critical low-dose region.

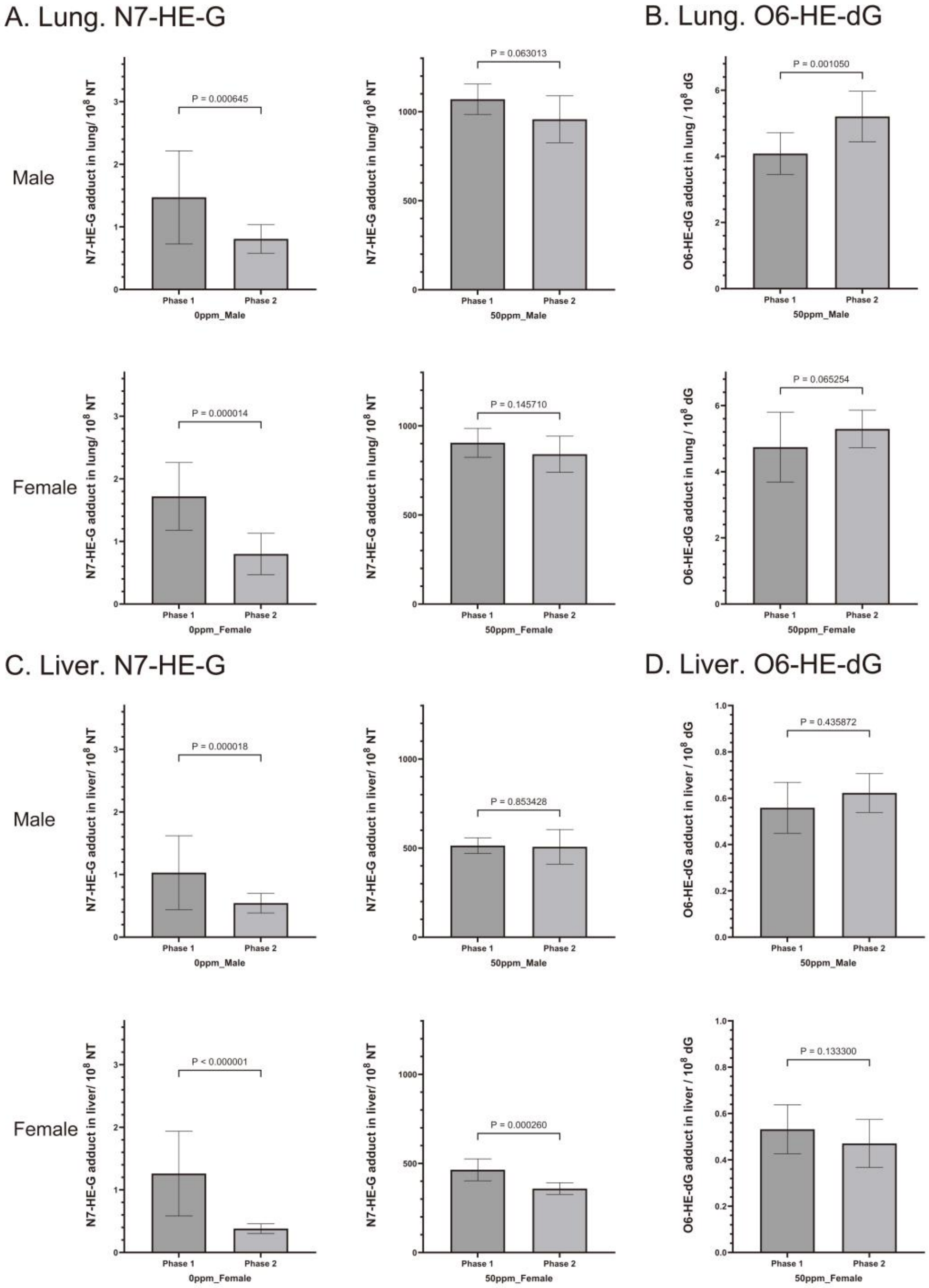


**Figure S1**. Comparative analysis of DNA adduct levels in the lung (A, B) and liver (C, D)

between experimental Phase 1 and Phase 2. To evaluate potential batch effects, levels of N7-HE-G (A, C) and $O^6$-HE-dG (B, D) were compared in the lung and liver of male and female mice from the shared exposure groups, 0 ppm (filtered air) and 50 ppm EtO.

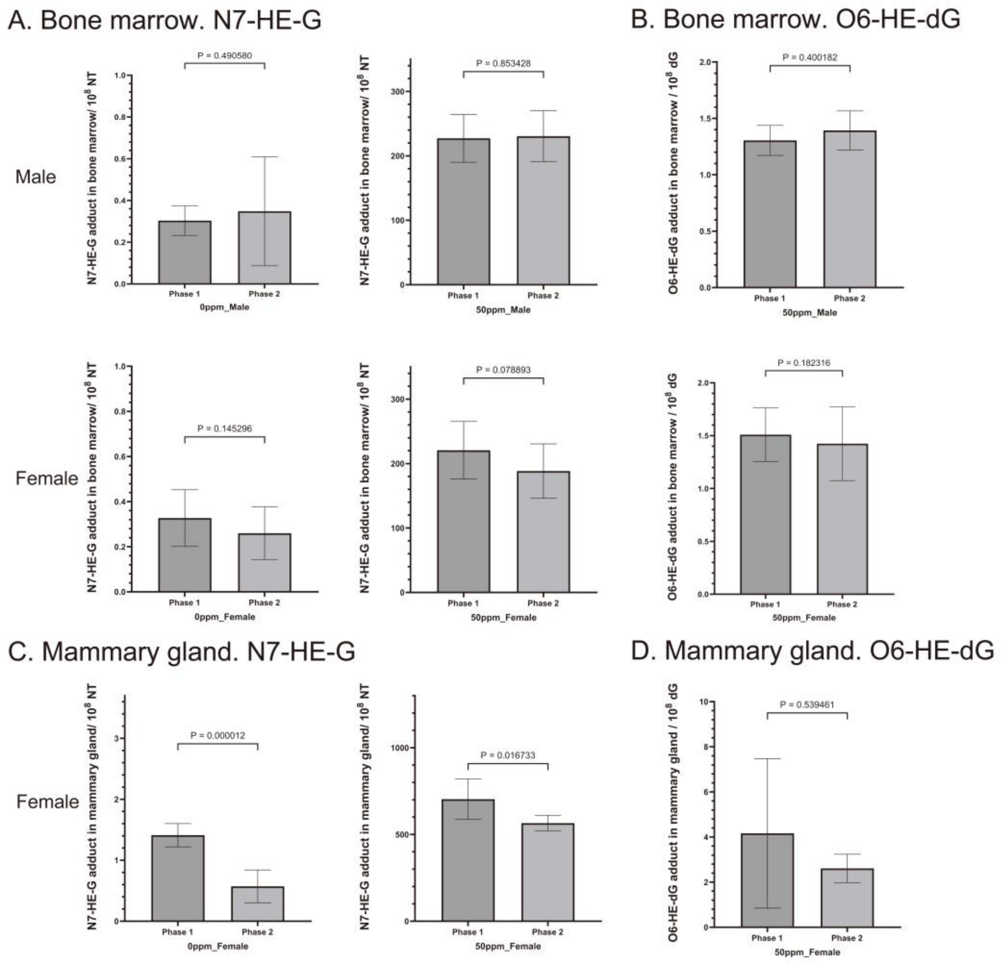


**Figure S2**. Comparative analysis of DNA adduct levels in the bone marrow (A, B) and mammary gland (C, D) between experimental Phase 1 and Phase 2. To evaluate potential batch effects, levels of N7-HE-G (A, C) and $O^6$-HE-dG (B, D) were compared in the bone marrow and mammary gland of mice from the shared exposure groups, 0 ppm (filtered air) and 50 ppm EtO.

### REFERENCES

Liu CW, Feng J, Peng J, Zhao H, Wang X, Gollapudi BB, Li AA, Bus JS, Kirman CR, Lu K. submitted. Molecular dosimetry of hemoglobin adducts in mice exposed to ethylene oxide. Archives of Toxicology. In revision.

Table S1. HPLC fractionation gradients for N7-HE-G and O6-HE-dG purification.

| For adduct | Time | %B* | Flow rate (mL/min) | Collection window** | Post time |
|---|---|---|---|---|---|
| N7-HE-G | 0 | 3.5 | 0.8 | 11.5 min to 13.5 min | 3 min |
| | 13 | 5 | 0.8 | | |
| | 22 | 22 | 0.8 | | |
| | 23 | 80 | 0.8 | | |
| | 27 | 80 | 0.8 | | |
| | 28 | 3.5 | 0.8 | | |
| | 33 | 3.5 | 0.8 | | |
| O6-HE-dG | 0 | 5 | 0.8 | 24 min to 26.1 min | 5 min |
| | 5 | 5 | 0.8 | | |
| | 12 | 15 | 0.8 | | |
| | 29 | 22 | 0.8 | | |
| | 30 | 80 | 0.8 | | |
| | 35 | 80 | 0.8 | | |
| | 36 | 5 | 0.8 | | |
| | 40 | 5 | 0.8 | | |

* A: 10 mM ammonium acetate in water; B: Methanol

** Collection windown may be slightly shifted using different batch of C18 column

Table S2. LC gradient conditions and MS parameters for LC-MS/MS analysis of N7-HE-G and O6-HE-dG.

| | | LC | | | | | MS | | PRM (Q Exactive HF) | | | | SRM (TSQ Quantis QqQ) | | | |
|---|---|---|---|---|---|---|---|---|---|---|---|---|---|---|---|---|
| For adduct | MS type | Time (min) | %B* | Flow rate (µL/min) | 10-port valve switch time (min) | Column temperature (°C) | Parameter | Setting | Precursor id | Precursor ion m/z | (N)CE | Target fragment ion m/z | Q1 m/z | Collision energy (V) | Q3 m/z | Dwell time (ms) |
| N7-HE-G | Q Exactive HF | 0 | 1 | 0.3 | | 35 | Polarity | Positive | N7-HE-G | 196.0829 | 40 | 152.0567 | | | | |
| | | 2.75 | 1 | 0.3 | 2.75 | | PRM mode | ON | N7-HE-G-d4 | 200.1080 | 40 | 152.0567 | | | | |
| | | 10 | 10 | 0.3 | | | Default charge state | 1 | | | | | | | | |
| | | 10.1 | 95 | 0.3 | | | Inclusion list | ON | | | | | | | | |
| | | 15 | 95 | 0.3 | | | MS2 microscans | 1 | | | | | | | | |
| | | 15.1 | 1 | 0.3 | | | MS2 resolution | 30000 | | | | | | | | |
| | | 24 | 1 | 0.3 | 18 | | AGC target | 1.00E+06 | | | | | | | | |
| | | | | | | | Maximum IT | 50 ms | | | | | | | | |
| | | | | | | | Loop count | 2 | | | | | | | | |
| | | | | | | | Isolation window (m/z) | 1.4 | | | | | | | | |
| N7-HE-G | TSQ Quantis QqQ | 0 | 1 | 200 | | 40 | Data mode | Centroid | N7-HE-G | | | | 196.1 | 19 | 152 | 300 |
| | | 0.3 | 1 | 200 | | | Collision gas pressure (mTorr) | 1.5 | N7-HE-G-d4 | | | | 200.1 | 19 | 152 | 300 |
| | | 3 | 10 | 200 | | | Q1 Resolution (FWHM) | 0.7 | | | | | | | | |
| | | 3.5 | 100 | 200 | | | Q3 Resolution (FWHM) | 0.7 | | | | | | | | |
| | | 6 | 100 | 200 | | | Polarity | Positive | | | | | | | | |
| | | 6.1 | 1 | 200 | | | | | | | | | | | | |
| | | 10 | 1 | 200 | | | | | | | | | | | | |
| O6-HE-dG | Q Exactive HF | 0 | 5 | 0.3 | | 35 | Polarity | Positive | O6-HE-dG | 312.1303 | 25 | 196.0828 | | | | |
| | | 3.75 | 5 | 0.3 | 3.75 | | PRM mode | ON | O6-HE-dG-d4 | 316.1554 | 25 | 200.1080 | | | | |
| | | 10 | 30 | 0.3 | | | Default charge state | 1 | | | | | | | | |
| | | 10.1 | 90 | 0.3 | | | Inclusion list | ON | | | | | | | | |
| | | 15 | 90 | 0.3 | | | MS2 microscans | 1 | | | | | | | | |
| | | 15.1 | 5 | 0.3 | | | MS2 resolution | 60000 | | | | | | | | |
| | | 22 | 5 | 0.3 | 18 | | AGC target | 5.00E+05 | | | | | | | | |
| | | | | | | | Maximum IT | 200 ms | | | | | | | | |
| | | | | | | | Loop count | 2 | | | | | | | | |
| | | | | | | | Isolation window (m/z) | 1.4 | | | | | | | | |

* A: 0.1% FA in water; B: 0.1% FA in ACN

Table S3. N7-HE-G adduct numbers (10^8 NT) quantified in the lung, liver, bone marrow, and mammary gland collected from mice exposed to EtO.

**Lung**

| | | Male | | | Female | | | |
|---|---|---|---|---|---|---|---|---|
| Group | Exposure concentration (ppm) | N7-HE-G | *n* | statistical significance between exposure group versus G1 | N7-HE-G | *n* | statistical significance between exposure group versus G1 | statistical significance between male and female in each exposure group |
| 1 | 0 | 1.250±0.693 | 30 | | 1.402±0.649 | 29 | | not significant |
| 2 | 0.05 | 1.957±0.402 | 10 | p < 0.001 | 1.640±0.551 | 10 | not significant | not significant |
| 3 | 0.1 | 3.160±0.731 | 10 | p < 0.001 | 2.524±0.745 | 10 | p < 0.001 | not significant |
| 4 | 0.5 | 17.827±3.534 | 10 | p < 0.001 | 17.772±4.141 | 10 | p < 0.001 | not significant |
| 5 | 1 | 30.539±2.412 | 10 | p < 0.001 | 27.984±5.148 | 10 | p < 0.001 | not significant |
| 6 | 50 | 1013.516±122.271 | 20 | p < 0.001 | 869.424±95.942 | 18 | p < 0.001 | p < 0.001 |
| 7 | 100 | 2317.856±329.587 | 10 | p < 0.001 | 2029.434±261.549 | 10 | p < 0.001 | p < 0.05 |
| 8 | 200 | 6434.504±871.137 | 10 | p < 0.001 | 6669.101±996.025 | 10 | p < 0.001 | not significant |

**Liver**

| | | Male | | | Female | | | |
|---|---|---|---|---|---|---|---|---|
| Group | Exposure concentration (ppm) | N7-HE-G | *n* | statistical significance between exposure group versus G1 | N7-HE-G | *n* | statistical significance between exposure group versus G1 | statistical significance between male and female in each exposure group |
| 1 | 0 | 0.868±0.539 | 30 | | 0.957±0.690 | 29 | | not significant |
| 2 | 0.05 | 1.277±0.242 | 10 | p < 0.005 | 0.863±0.187 | 10 | not significant | p < 0.001 |
| 3 | 0.1 | 2.183±0.803 | 9 | p < 0.001 | 1.251±0.190 | 10 | p < 0.05 | p < 0.01 |
| 4 | 0.5 | 8.206±1.691 | 9 | p < 0.001 | 8.006±1.063 | 10 | p < 0.001 | not significant |
| 5 | 1 | 14.923±2.239 | 10 | p < 0.001 | 13.568±2.361 | 10 | p < 0.001 | not significant |
| 6 | 50 | 510.919±73.447 | 20 | p < 0.001 | 408.950±71.587 | 19 | p < 0.001 | p < 0.001 |
| 7 | 100 | 1238.133±267.363 | 10 | p < 0.001 | 1137.923±175.998 | 10 | p < 0.001 | not significant |
| 8 | 200 | 4449.120±833.577 | 10 | p < 0.001 | 4796.690±1044.021 | 10 | p < 0.001 | not significant |

**Bone marrow**

| | | Male | | | Female | | | |
|---|---|---|---|---|---|---|---|---|
| Group | Exposure concentration (ppm) | N7-HE-G | *n* | statistical significance between exposure group versus G1 | N7-HE-G | *n* | statistical significance between exposure group versus G1 | statistical significance between male and female in each exposure group |
| 1 | 0 | 0.319±0.160 | 29 | | 0.305±0.125 | 30 | | not significant |
| 2 | 0.05 | 0.624±0.164 | 10 | p < 0.001 | 0.596±0.276 | 10 | p < 0.01 | not significant |
| 3 | 0.1 | 0.907±0.263 | 10 | p < 0.001 | 0.643±0.087 | 9 | p < 0.001 | p < 0.05 |
| 4 | 0.5 | 3.397±0.698 | 8 | p < 0.001 | 3.222±0.688 | 10 | p < 0.001 | not significant |
| 5 | 1 | 5.581±0.988 | 10 | p < 0.001 | 5.924±1.825 | 10 | p < 0.001 | not significant |
| 6 | 50 | 228.798±37.355 | 20 | p < 0.001 | 203.689±45.282 | 19 | p < 0.001 | not significant |
| 7 | 100 | 503.278±100.342 | 10 | p < 0.001 | 453.734±70.618 | 10 | p < 0.001 | not significant |
| 8 | 200 | 1348.527±276.057 | 10 | p < 0.001 | 1650.603±443.751 | 10 | p < 0.001 | not significant |

**Mammary gland**

| | | Female | | |
|---|---|---|---|---|
| Group | Exposure concentration (ppm) | N7-HE-G | *n* | statistical significance between exposure group versus G1 |
| 1 | 0 | 1.030±0.483 | 22 | |
| 2 | 0.05 | 1.238±0.148 | 10 | not significant |
| 3 | 0.1 | 1.994±0.178 | 10 | p < 0.001 |
| 4 | 0.5 | 13.881±2.880 | 5 | p < 0.001 |
| 5 | 1 | 21.635±3.377 | 6 | p < 0.001 |
| 6 | 50 | 616.978±102.874 | 16 | p < 0.001 |
| 7 | 100 | 1445.920±268.381 | 6 | p < 0.001 |
| 8 | 200 | 5806.243±1063.218 | 5 | p < 0.001 |

Table S4. O6-HE-dG adduct numbers (10^8 dG) quantified in the lung, liver, bone marrow, and mammary gland collected from mice exposed to EtO.

| | | Lung | | | | | Liver | | | | | Bone marrow | | | | | Mammary gland | |
|---|---|---|---|---|---|---|---|---|---|---|---|---|---|---|---|---|---|---|
| | | Male | | Female | | | Male | | Female | | | Male | | Female | | | Female | |
| Group | Exposure concentration (ppm) | O6-HE-dG | *n* | O6-HE-dG | *n* | statistical significance between male and female in each exposure group | O6-HE-dG | *n* | O6-HE-dG | *n* | statistical significance between male and female in each exposure group | O6-HE-dG | *n* | O6-HE-dG | *n* | statistical significance between male and female in each exposure group | O6-HE-dG | *n* |
| 1 | 0 | ND* | 30 | ND | 30 | | ND | 30 | ND | 30 | | ND | 24 | ND | 23 | | ND | 15 |
| 2 | 0.05 | ND | 10 | ND | 10 | | ND | 10 | ND | 10 | | ND | 10 | ND | 7 | | ND | 8 |
| 3 | 0.1 | ND | 10 | ND | 10 | | ND | 10 | ND | 10 | | ND | 10 | ND | 9 | | ND | 10 |
| 4 | 0.5 | ND | 10 | ND | 10 | | ND | 10 | ND | 10 | | ND | 8 | ND | 10 | | ND | 4 |
| 5 | 1 | ND | 10 | ND | 10 | | ND | 10 | ND | 10 | | ND | 10 | ND | 9 | | ND | 4 |
| 6 | 50 | 4.646±0.891 | 20 | 5.033±0.859 | 19 | not significant | 0.591±0.101 | 20 | 0.500±0.106 | 19 | $p < 0.01$ | 1.351±0.158 | 19 | 1.465±0.303 | 19 | not significant | 3.052±1.828 | 14 |
| 7 | 100 | 9.992±2.253 | 10 | 11.549±1.806 | 10 | not significant | 1.260±0.402 | 10 | 1.197±0.277 | 10 | not significant | 2.869±0.448 | 10 | 3.101±0.443 | 10 | not significant | 8.503±4.898 | 4 |
| 8 | 200 | 33.750±6.021 | 10 | 37.004±7.061 | 10 | not significant | 4.410±0.865 | 10 | 4.710±1.106 | 10 | not significant | 9.000±1.296 | 10 | 10.512±3.169 | 10 | not significant | 26.501±9.023 | 4 |

* ND: not detected
